# Preliminary attempt to inhibit proliferation of NUT carcinoma cell lines using antisense oligonucleotides targeting NUTM1

**DOI:** 10.1101/2024.07.30.605673

**Authors:** Jessica Alley, Benjamin G. Vincent, Alexander Rubinsteyn

## Abstract

NUT carcinoma is an aggressive cancer driven by NUTM1 fusion proteins. This study evaluated antisense oligonucleotides (ASOs) targeting NUTM1 as a potential therapeutic approach. Three ASOs targeting NUTM1 and scrambled controls were tested at 10-50 nM doses in three NUT carcinoma cell lines (TC-797, 10-15, 14169) and one control line (293T). ASOs were delivered both gymnotically and using a transfection reagent. Cell viability was assessed at 48 and 72 hours using a luminescence-based assay. No ASOs showed selective inhibition of NUT carcinoma cell viability compared to controls across all conditions tested. Thus, this pilot study did not identify ASOs with activity against NUT carcinoma cells. However, it did establish preliminary protocols and generated data to inform future ASO optimization efforts targeting NUTM1 in NUT carcinoma. Further refinement of ASO design and delivery methods is needed to identify therapeutic candidates.

**Graphical Abstract:** 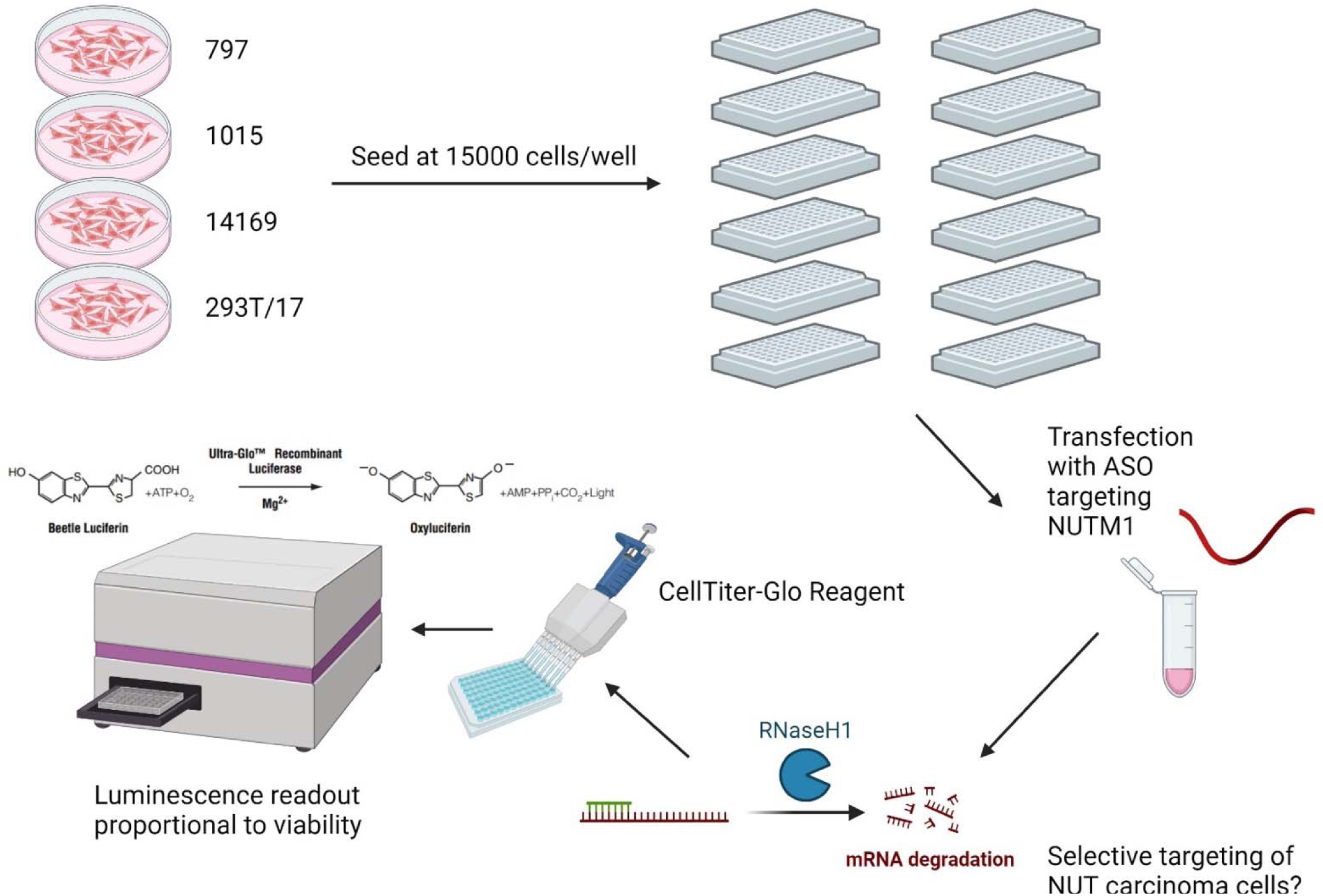

**Hypothesis:** This experiment was performed as a pilot experiment to generate data to inform future work, which should include formal hypothesis testing with a reduced number of conditions. The hypothesis of this pilot experiment was that one or more of the non-scrambled antisense oligonucleotides (ASOs) targeting NUTM1 will selectively decrease proliferation and/or viability of NUT carcinoma cells, indicated by a reduction in luminescence (as compared to treated non-NUT carcinoma and untreated NUT carcinoma control cells) using the CellTiter-Glo Luminescent Cell Viability Assay.

## Background and Rationale

NUT carcinoma is a highly aggressive cancer currently lacking effective treatments (French et al., 2022). Because it is driven by the BRD4-NUT fusion oncoprotein in approximately 75% of cases (Eagen & French, 2021), one novel therapeutic strategy could include targeting translation of this fusion protein through ASOs, which are short (usually between 12 and 30 nucleotides) synthetic nucleic acids designed to base pair with cognate RNA and modulate its function through diverse mechanisms, including targeting the RNA for degradation or alternative splicing, dependent upon the specificity of the ASO (Bennett et al., 2019). However, by disrupting inhibitory elements, ASOs can also be designed to enhance translation (Liang et al., 2017). Thus, validation of the intended effects of ASOs through initial *in vitro* experiments is critical.

For this work, three ASOs have been designed to target the coding sequence of NUTM1 (from exon 3 through part of exon 8) via *in silico* selection for dissimilarity from other genes as optimizing for heuristic criteria such as shorter repeat stretches (Table 1). These ASOs are each 21 nucleotides in length and target sequences of NUTM1 shared by three human NUT carcinoma cell lines (797, 1015, and 14169). To limit potential off-target toxicity, these candidates do not share 14mer subsequences with any other transcripts in mice or humans. Three scrambled ASOs (containing the same respective nucleic acids in a randomized order) were tested as controls alongside their non-scrambled counterparts. The use of these scrambled ASOs provided information regarding the inherent toxicity of ASOs, independent of sequence, which was another motivation for this work. Of the scrambled ASOs, two share 15mer subsequences and one shares 14mer subsequences with other (non- NUTM1) transcripts. All six ASOs include a backbone with phosphorothioate bonds, which prolongs degradation by nucleases and clearance from the blood, enhancing tissue uptake *in vivo* (Geary et al., 2015).

**Table 1.**
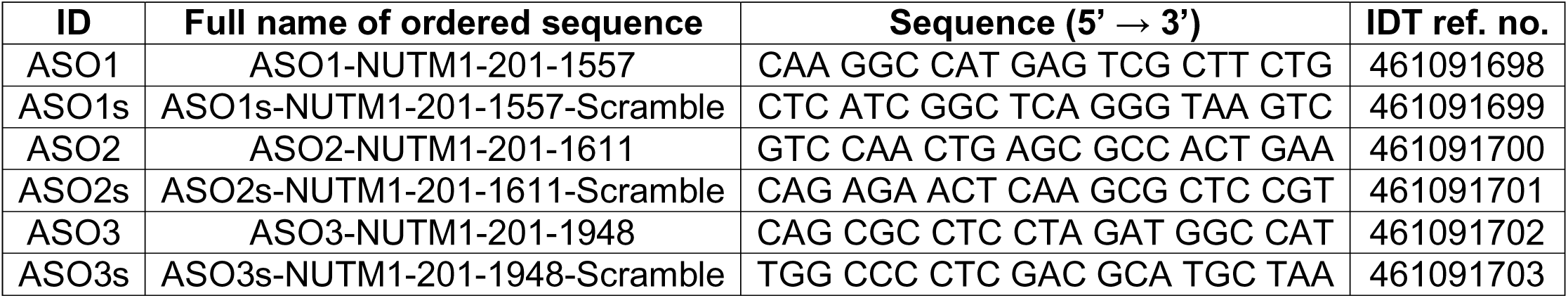
Description of ASOs tested in this experiment.

These ASOs were designed with the goal of reducing expression of the BRD4-NUT fusion protein through targeting of cognate mRNA transcripts for degradation by endogenous nucleases such as ribonuclease H1 that recognize and cleave DNA-RNA duplexes. Because the BRD4-NUT oncoprotein is thought to be the sole driver of malignant transformation in NUT carcinoma (Durall et al., 2023) and its knockdown has been shown to result in cell differentiation and growth arrest (French et al., 2008), luminescence proportional to viable cell number using Promega’s CellTiter-Glo Luminescent Cell Viability Assay (kit G7572) was used as the readout to evaluate efficacy of the ASOs tested in this experiment. In this assay system, a single reagent is added to serum-containing culture medium. The assay reagent lyses cells and generates a luminescent signal proportional to the amount of ATP present, the source of which is viable cells (Figure 1).

**Figure 1.**
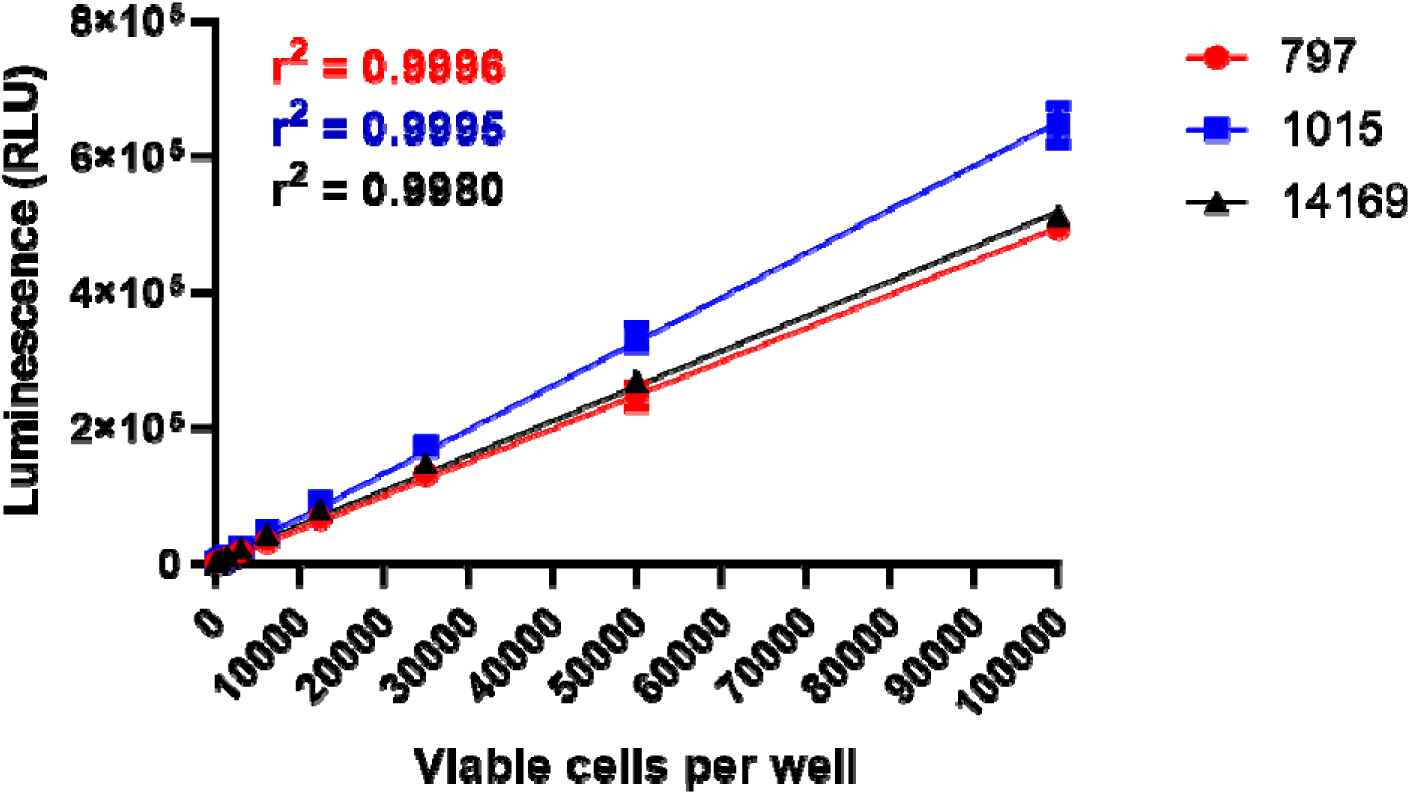
NUT carcinoma cell number correlates with luminescent output using Promega kit G7572. NUT carcinoma cell lines were plated in white 96-well plates at the concentrations indicated using 1:2 serial dilutions in culture medium. Approximately 30 min later, an equal volume (100 µL) of Promega CellTiter-Glo Reagent was added followed by shaking at 300 rpm for 5 min. Luminescence was recorded after an additional 5 min. Data are presented as means of three replicates for each measurement, with error bars indicating standard deviations and lines of best fit shown.

An alternative experimental approach could include assessment of the luciferase-expressing NUT carcinoma cell line counterparts using methods optimized during pilot experiments (Figures 2-4). However, the requirement for a single reagent in Promega kit G7572 renders this method higher throughput and ideal for large experiments such as this pilot. Additionally, the kit is versatile, working with both adherent or suspension cells and when added to either DPBS or serum-containing media, and having sufficient reagent already available in-house makes using Promega kit G7572 the more economical approach.

**Figure 2.**
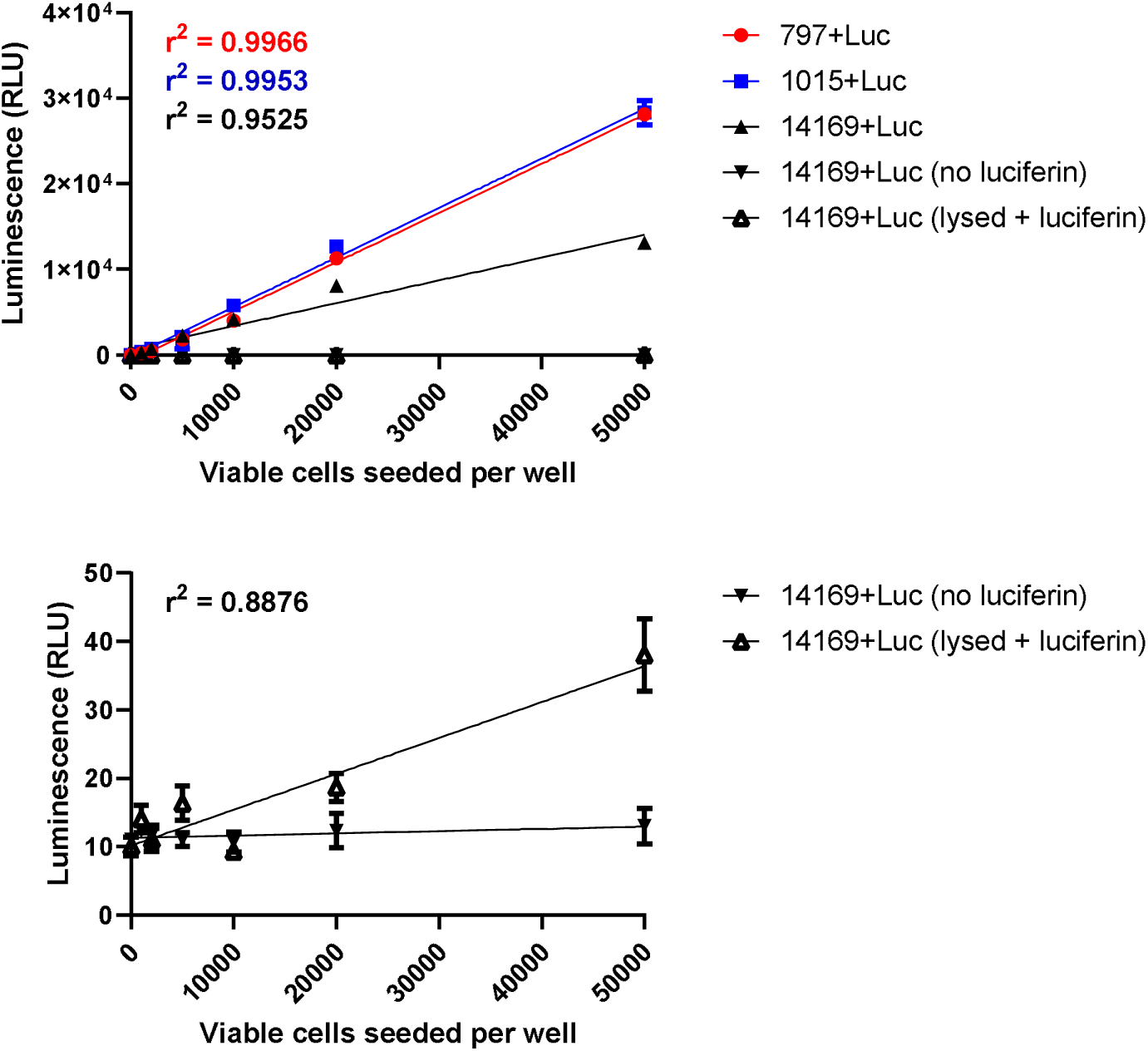
Luciferase-expressing NUT carcinoma cell number correlates with luminescent output using luciferin, although lysing cells and transferring lysates to assay plate reduces linear signal. Luciferase-expressing NUT carcinoma cell lines were seeded in 96-well plates at the concentrations indicated. After two days, media was removed and wells were washed with 200 µL DPBS. To lyse cells, 80 µL lysis reagent (prepared from Promega kit E1500) was added followed by shaking at 300 rpm for 5 min. Lysates were transferred to a white plate and 20 µL luciferin (Promega P1041) was added such that final well concentration was 250 µg/mL. For all other conditions, 100 µL DPBS (no luciferin) or 100 µL luciferin diluted to 250 µg/mL in DPBS was added. Luminescence was recorded 10 min after the addition of luciferin. Data are presented as means of three replicates for each measurement, with error bars indicating standard deviations and lines of best fit shown. Bottom panel depicts data found in top panel, with adjusted y-axis and r-squared value corresponding to lysed cells.

**Figure 3.**
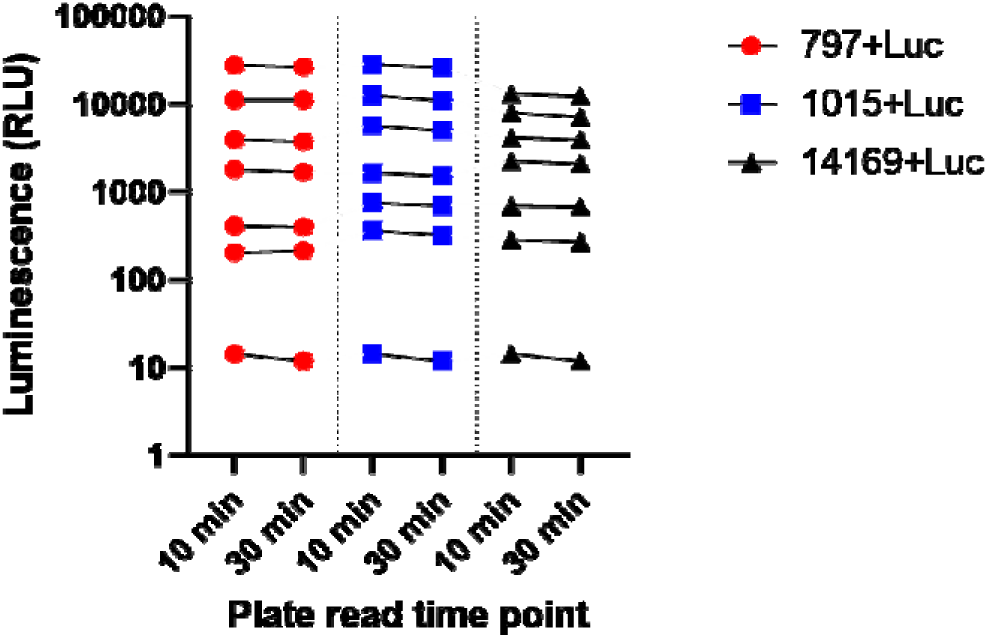
Luminescent output of luciferase-expressing NUT carcinoma cells can be measured 10 min after the addition of luciferin. Luciferase-expressing NUT carcinoma cell lines were seeded in 96-well plates at concentrations ranging from 1000 to 50000 viable cells per well. After two days, media was removed and wells were washed with 200 µL DPBS. 100 µL luciferin (Promega P1041) diluted to 250 µg/mL in DPBS was added, and luminescence was recorded after 10 and 30 min. Data are presented as means of three replicates for each measurement.

**Figure 4.**
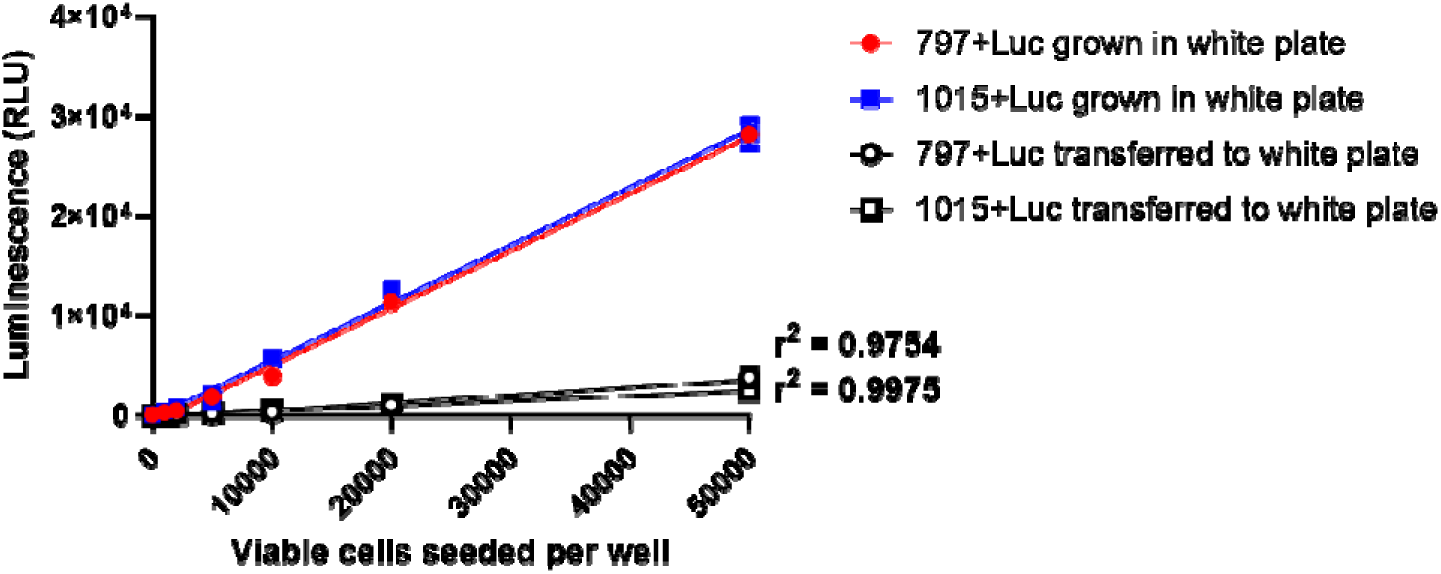
Transferring contents of unlysed wells containing luciferin reduces luminescent signal but not linear relationship with luciferase-expressing cell number. Luciferase-expressing NUT carcinoma cell lines were seeded in 96-well plates at the concentrations indicated. After two days, media was removed and wells were washed with 200 µL DPBS. For cells grown in white plates, 100 µL luciferin (Promega P1041) diluted to 250 µg/mL in DPBS was added, and luminescence was recorded immediately. For cells not grown in white plates, luciferin was similarly added then well contents were transferred to a white plate immediately prior to recording luminescence. Data are presented as means of three replicates for each measurement, with error bars indicating standard deviations and lines of best fit shown.

While some cell types can internalize ASOs through endocytosis or micropinocytosis, it is currently unknown if NUT carcinoma cells are capable of this. In parallel with gymnotic delivery, the use of a transfection reagent should facilitate entry of the ASOs into the cells and enable this experiment to be higher throughput and more cost effective than electroporation. Thus, in the current experiment, RNAiMAx was used given that it has previously yielded high transfection efficiency with low cytotoxicity as compared to other transfection reagents, especially Lipofectamine 2000, when used to deliver ASOs to multiple cell types (Wang et al., 2018) and been found to successfully deliver mRNA to these NUT carcinoma cell lines with limited cytotoxicity in pilot experiments (Figures 5 and 6).

**Figure 5.**
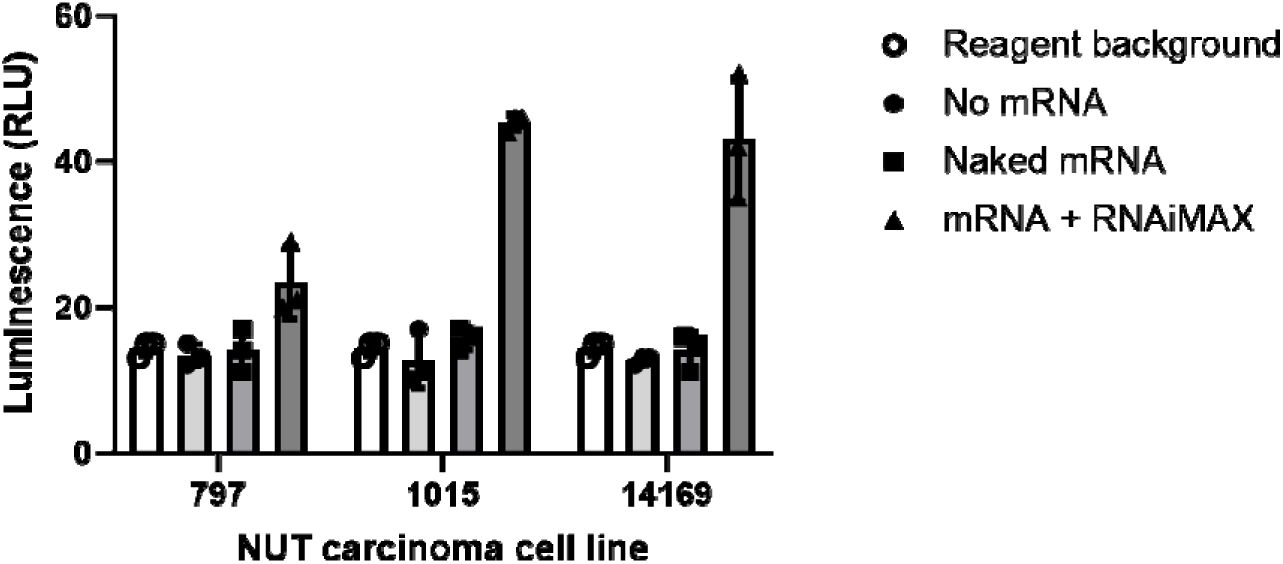
Recommended RNAiMAX dose enables transfection of NUT carcinoma cells with mRNA encoding luciferase. NUT carcinoma cell lines were seeded in 96-well plates at 50000 viable cells per well. The next day, when cells were 50-80% confluent, 100 ng mRNA encoding firefly luciferase was added. Culture wells of 200 µL serum- containing media received either 10 uL of mRNA diluted in Opti-MEM (Naked mRNA) or mixed with Lipofectamine RNAiMAX transfection reagent (mRNA + RNAiMAX) following manufacturer’s protocol. After 24 h, media was removed and wells were washed with 200 µL DPBS. 100 µL luciferin (Promega P1041) diluted to 250 µg/mL in DPBS was added, and well contents were transferred to a white plate immediately prior to recording luminescence. Reagent background wells containing only luciferin (no cells) were also included. Data are presented as means of three replicates for each measurement with error bars indicating standard deviations.

**Figure 6.**
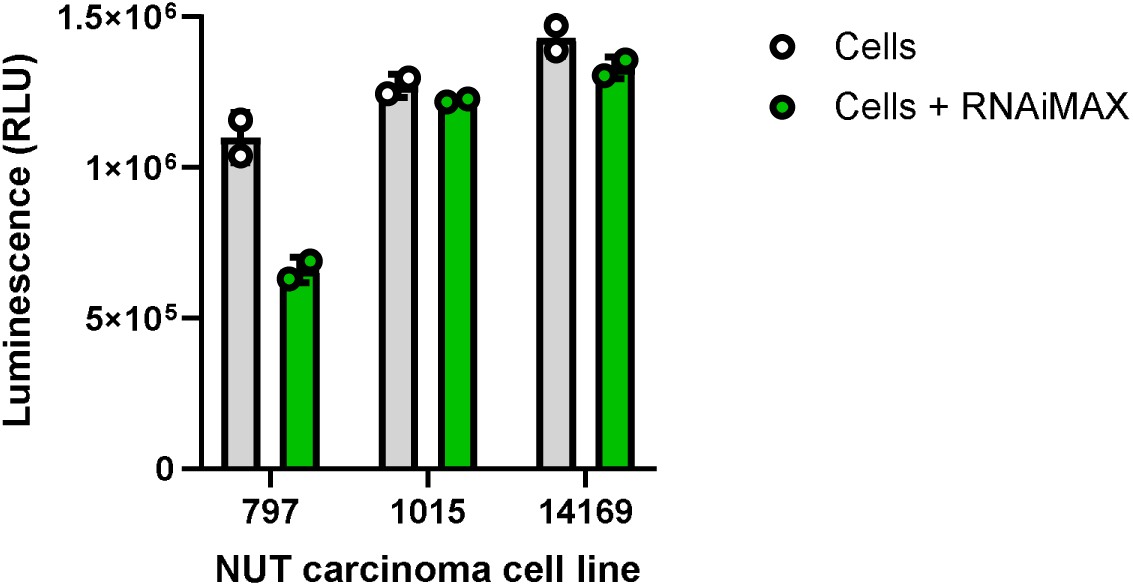
The 797 NUT carcinoma cell line may be more sensitive to RNAiMAX toxicity, or toxicity of this transfection reagent may be mediated by cell density at the time that it is introduced. NUT carcinoma cell lines were seeded in 96-well plates at 20000 viable cells per well. The next day, indicated wells (Cells + RNAiMAX) of cells in 200 µL serum-containing media received 10 µL of Opti-MEM plus Lipofectamine RNAiMAX transfection reagent prepared following manufacturer’s protocol. At this time point, 797 wells were approximately 30% confluent while all other wells appeared approximately 50% confluent. After 72 h of incubating with or without the transfection reagent, media was replaced with 100 µL DPBS. An equal volume of Promega CellTiter-Glo Reagent (from kit G7572) was added followed by shaking at 300 rpm for 5 min. Well contents were then transferred to a white plate, and luminescence was recorded after an additional 5 min. Data are presented as means of two replicates for each measurement with error bars indicating standard deviations.

Following completion of these experiments, ASOs that selectively decrease NUT carcinoma cell viability or hinder proliferation may be further pursued, potentially including additional evaluation of off-target toxicity such as assessment of hepatoxic potential (Sewing et al., 2016) or other recommended predictive assays (Goyenvalle et al., 2023). Next steps could also include adjustments to ASOs that show promise but require a transfection reagent to enter cells (such as shortening sequence length or conjugating to antibodies or aptamers that target receptors found on the surface of NUT carcinoma cells, like B7-H3) along with sugar modifications to decrease immunogenicity (such as 2’-MOE) or design considerations to increase stability (such as locked nucleic acid gapmers).

Pending progress of related mouse work, efficacy of candidate ASOs could later be tested in NUT carcinoma xenografts in immunodeficient mice.

### Design Summary

Three human NUT carcinoma cell lines (797, 1015, and 14169) and one non-NUT carcinoma cell line (293T/17) were propagated in DMEM supplemented with FBS, penicillin-streptomycin, and L-glutamine until adequate cell numbers (at least 5 x 10^6 cells per cell line) were generated. Cells were then seeded at 15000 viable cells per well in white plates and transfected with ASOs the following day. Transfection using the Lipofectamine RNAiMAX transfection reagent following the manufacturer’s protocol occurred alongside gymnotic delivery of ASOs. Cell viability was assessed using the CellTiter-Glo Luminescent Cell Viability Assay 48 and 72 h after transfection. In pilot work, this assay resulted in spillover from brightest wells into adjacent wells below 0.1% of signal in white plates, thus plate layouts in the current experiment should not need to be designed to mitigate this.

While not a perfect comparison, the time points were selected based on Schwartz et al. (2011), in which viability of NUT carcinoma cells was reduced to approximately 40% of controls after three days of growth in the HDAC inhibitor vorinostat. Based on pilot work, this time frame should allow introduction of the transfection reagent when cells are approximately 30-50% confluent (which corresponds to the data used to generate Figure 6, above) and completion of the experiment near the point when untreated cells will become overly confluent and turn the media acidic, which could confound viability outcomes.

Three ASO doses were tested in this experiment (10, 25, and 50 nM). These doses are within the range used *in vitro* in previous ASO studies (Liang et al., 2017; Zhang et al., 2023) and overlap with the range of siRNA doses used by Jeff Jensen in his recent experiments with the same NUT carcinoma cell lines as well as that utilized in previously published work by Grayson et al. (2014) using siRNA targeting NUTM1 in NUT carcinoma cells. Each ASO dose was tested in triplicate in this experiment, yielding a total of at least 912 wells, including controls:

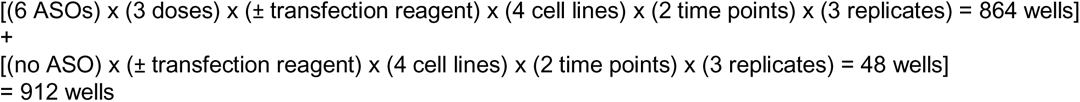

Conditions tested included the following for each cell line, with plate layouts below:

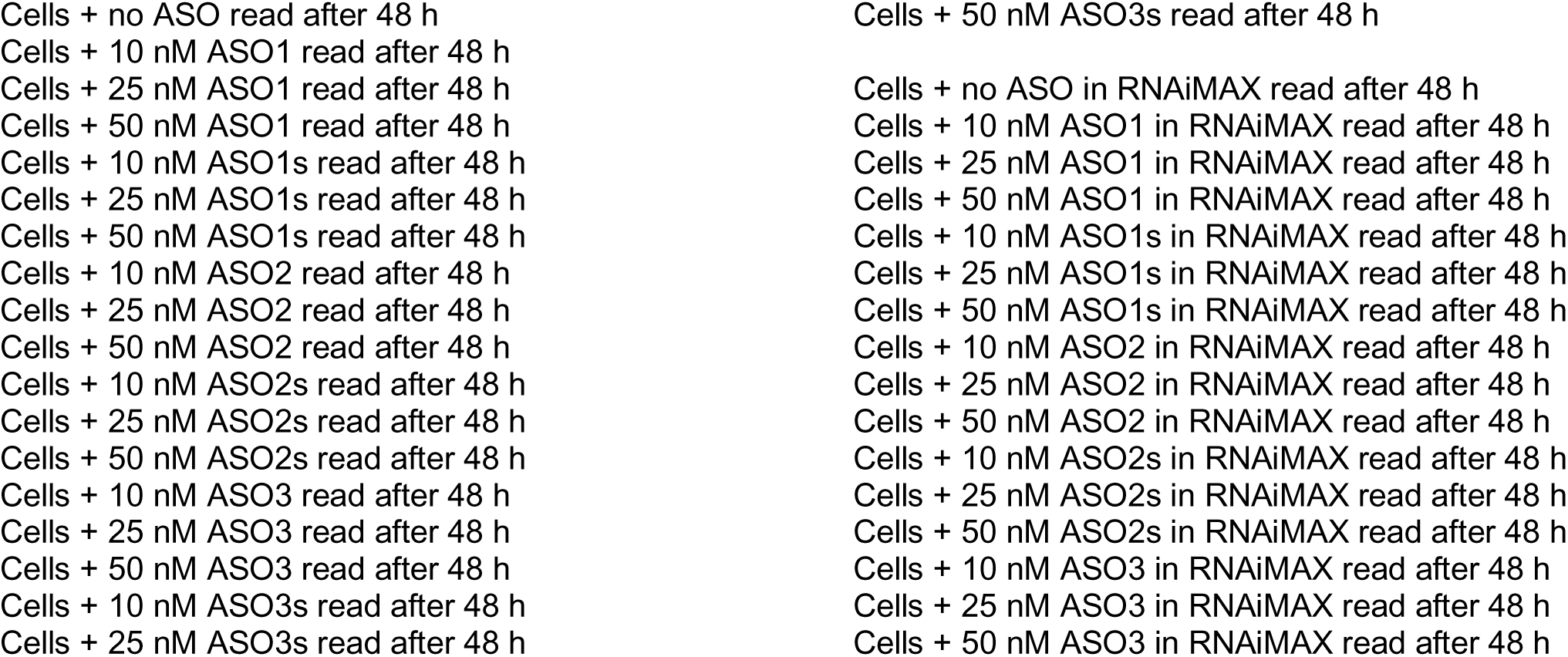
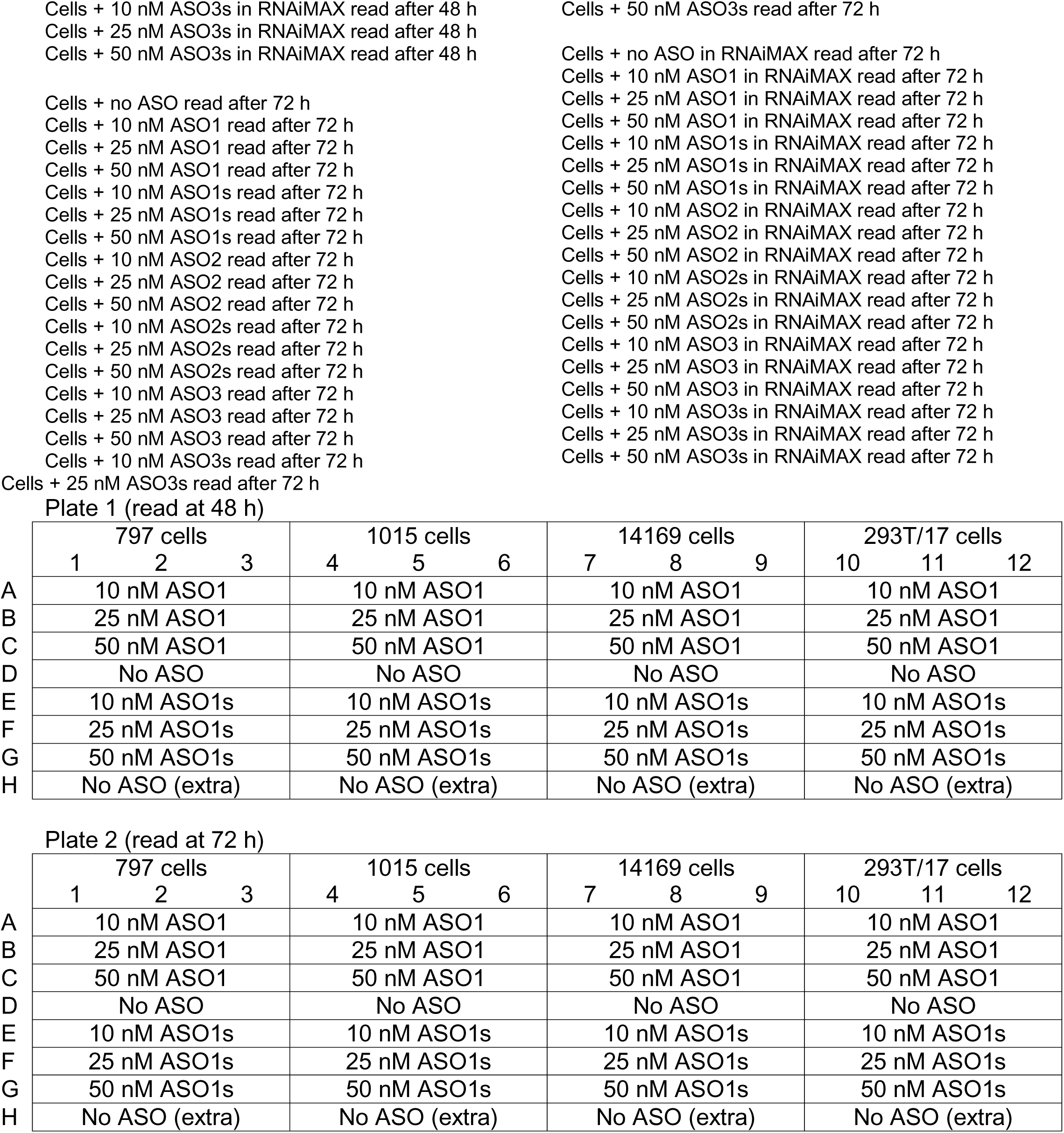
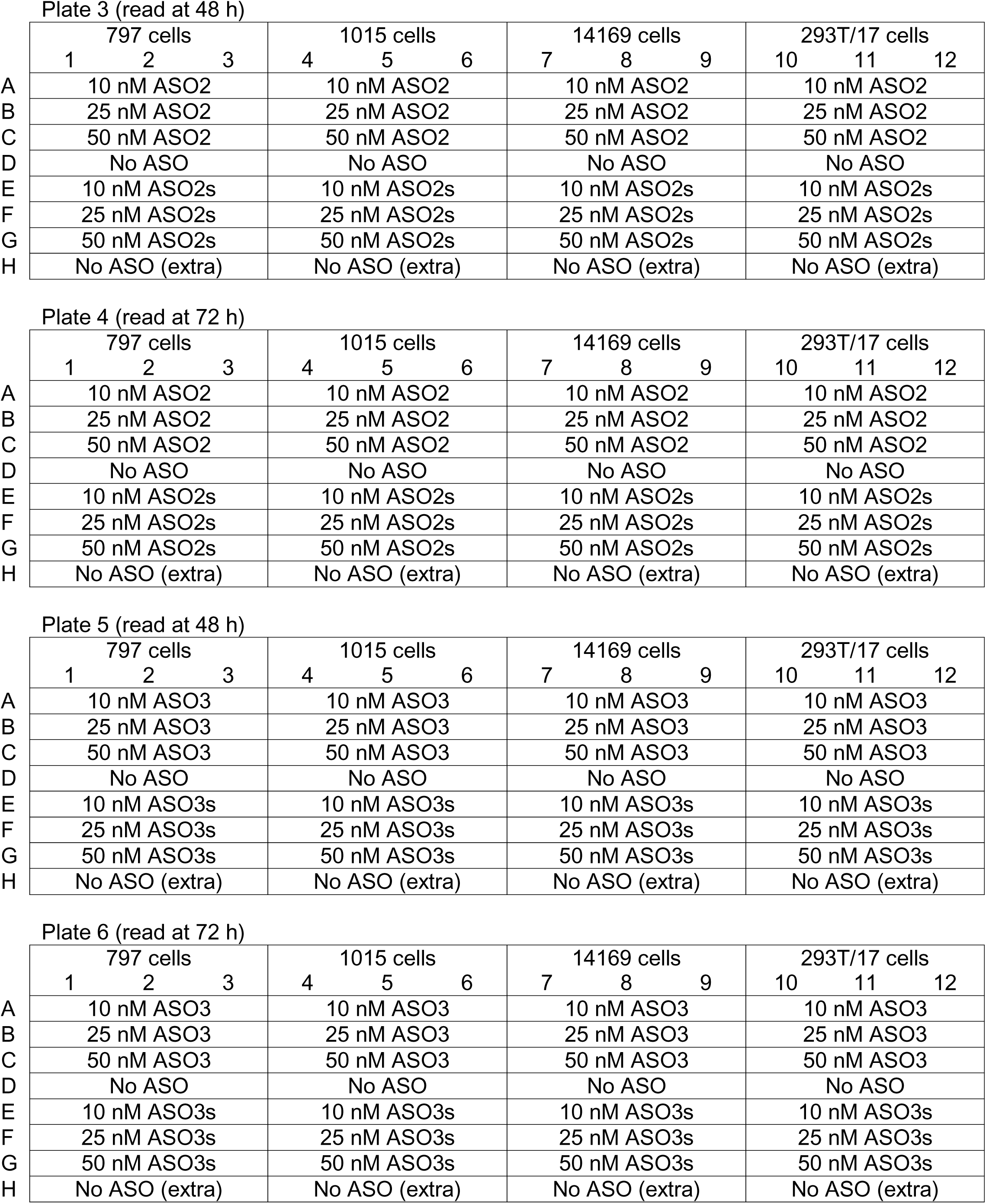
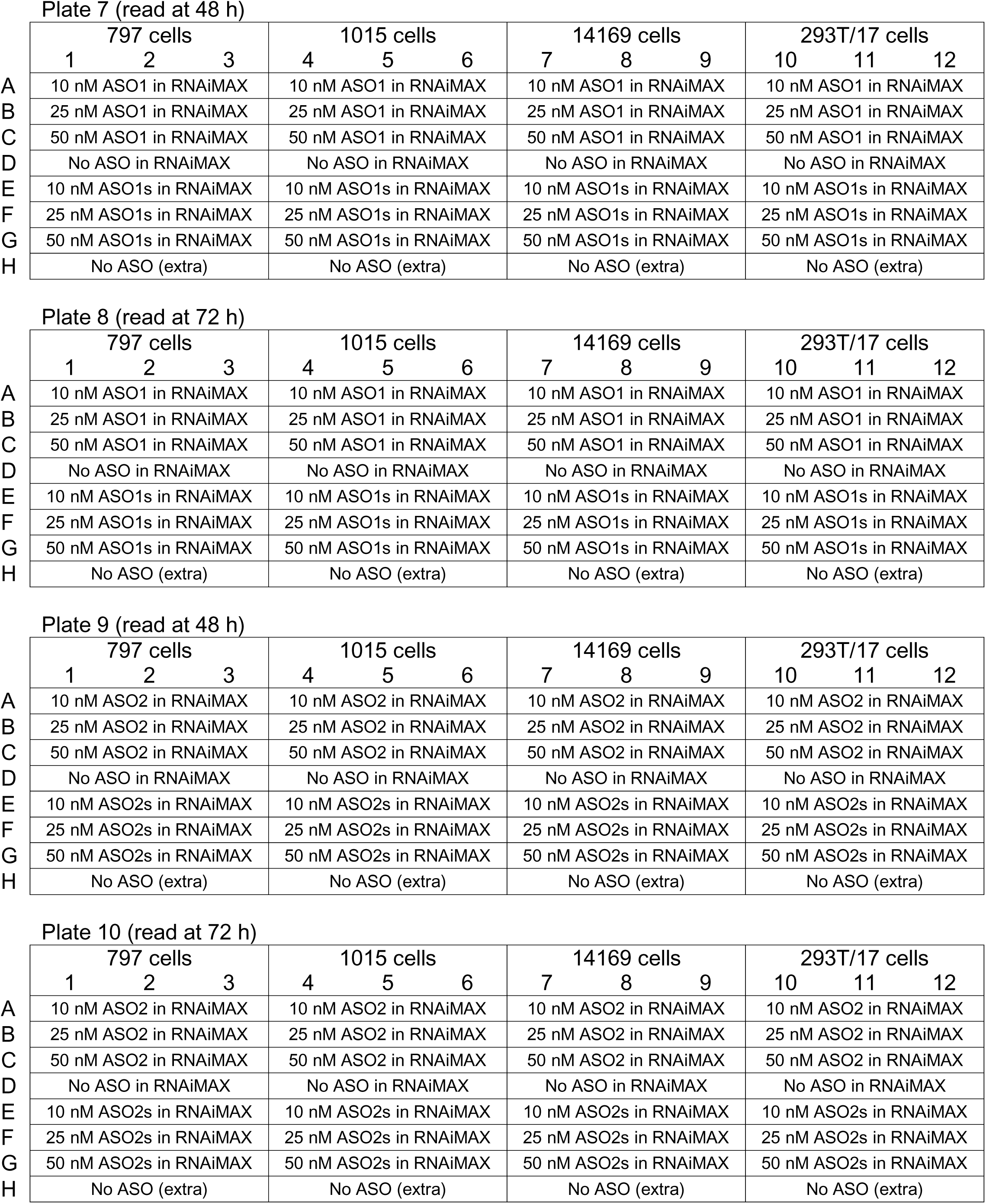
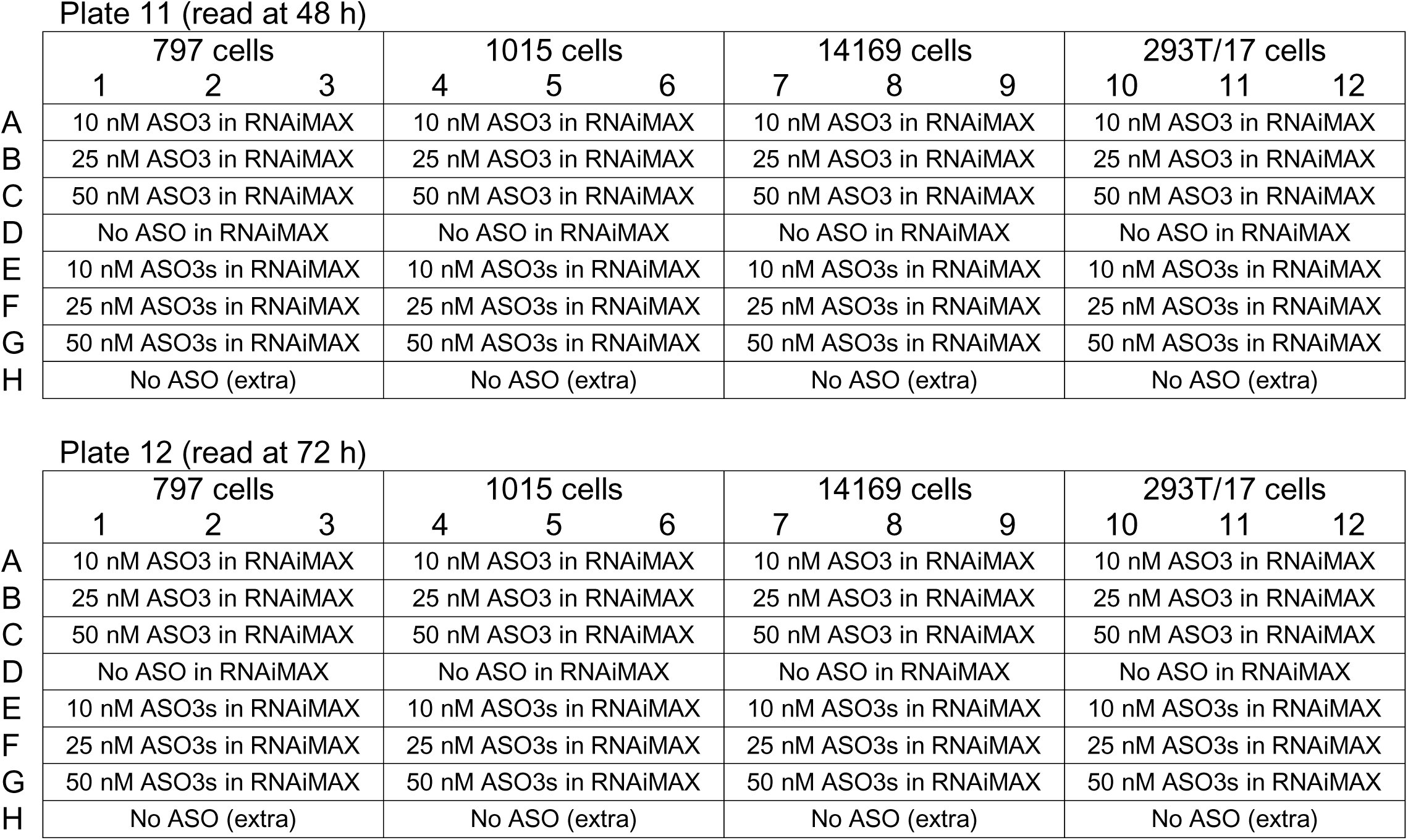

### Analysis Plan and Statistical Considerations

Luminescence was recorded for all wells using the Synergy 2 plate reader, and triplicates were then averaged for each condition. Because there were cells in every well of each plate, no reagent background wells were included for subtraction from condition/group means. However, in pilot experiments, background values were low (below 25 RLU), so this was not anticipated to be a concern since values for all wells in the current experiment were expected to be multiple orders of magnitude higher.

While this pilot experiment was not designed for statistical power, GraphPad Prism was used to generate graphs to identify patterns across conditions. These data will be used to determine next steps with the ASO work as described in the Background and Rationale section above. Specifically, any ASO that selectively decreases NUT carcinoma cell viability or hinders proliferation (identified as reduced luminescence) as compared to controls may be further pursued in a hypothesis-testing study using effect size and variance estimated from the current experiment, and methods or ASO design may be further optimized.

## Results

Neither oligonucleotides delivered gymnotically (Figures 7 & 8) nor oligonucleotides delivered with Lipofectamine RNAiMAX (Figures 9 & 10) showed a meaningful difference between oligonucleotides targeting NUTM1 (ASO1, ASO2, ASO3) and their scrambled equivalents (ASO1s, ASO2s, ASO3s). This was true both in terms of raw luminescence values (Figures 7 & 9) as well as relative to negative control (Figures 8 & 10).

**Figure 7.**
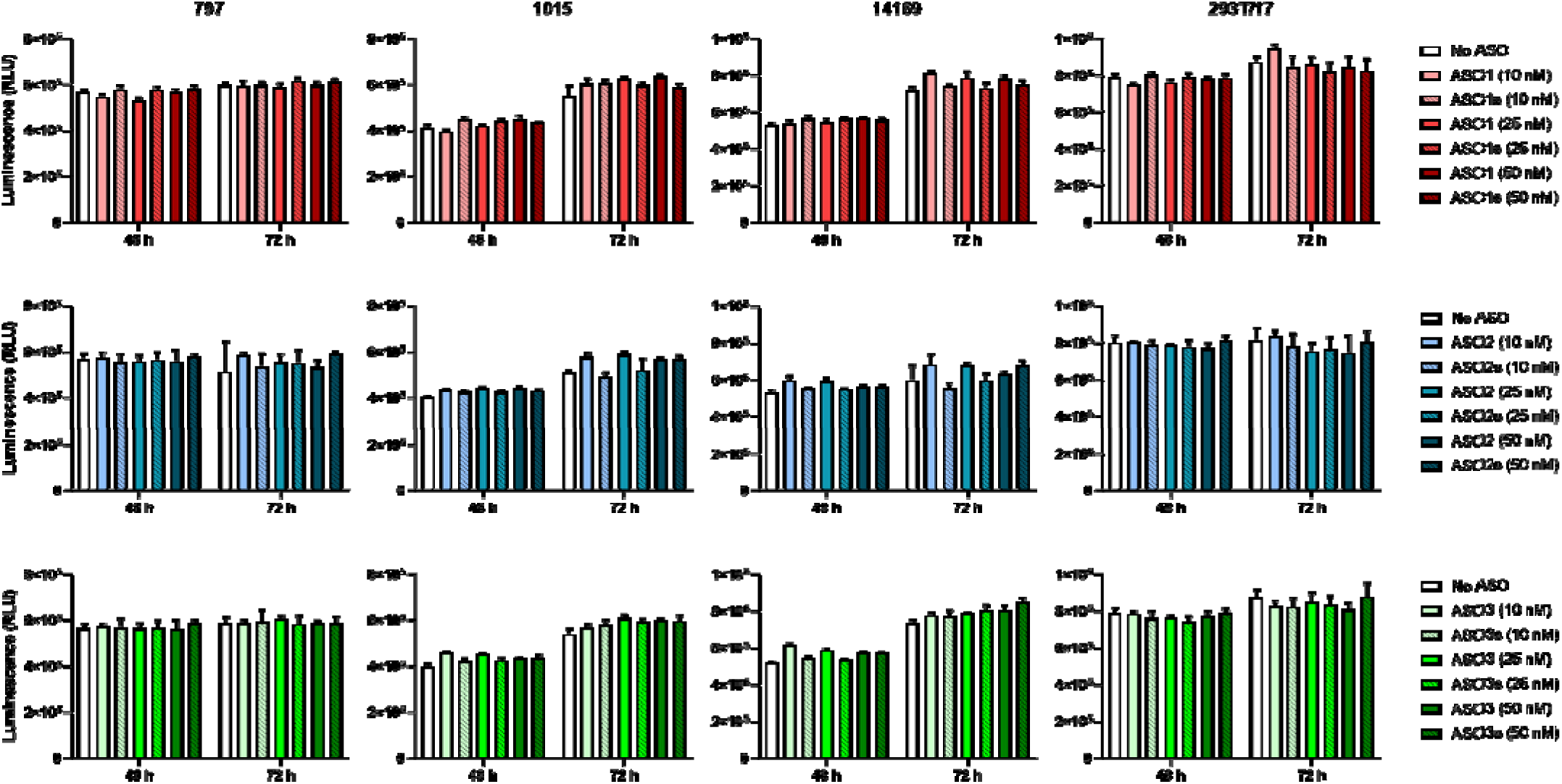
Three NUT carcinoma (797, 1015, 14169) and one non-NUT carcinoma (293T/17) cell lines were seeded in white 96-well plates at 15000 viable cells/well. The next day, antisense oligonucleotides targeting NUTM1 (ASO1, ASO2, ASO3) or scrambled counterparts containing the same respective nucleic acids in randomized order (ASO1s, ASO2s, ASO3s) were delivered in 10 µL Opti-MEM to reach the doses indicated. Either 48 or 72 h later, culture medium was replaced with DPBS and an equal volume (100 µL) of Promega CellTiter-Glo Reagent was added followed by shaking at 300 rpm for 5 min. Luminescence was recorded after an additional 5 min. Data are presented as means of three or six replicates for each measurement, with error bars indicating standard deviations.

**Figure 8.**
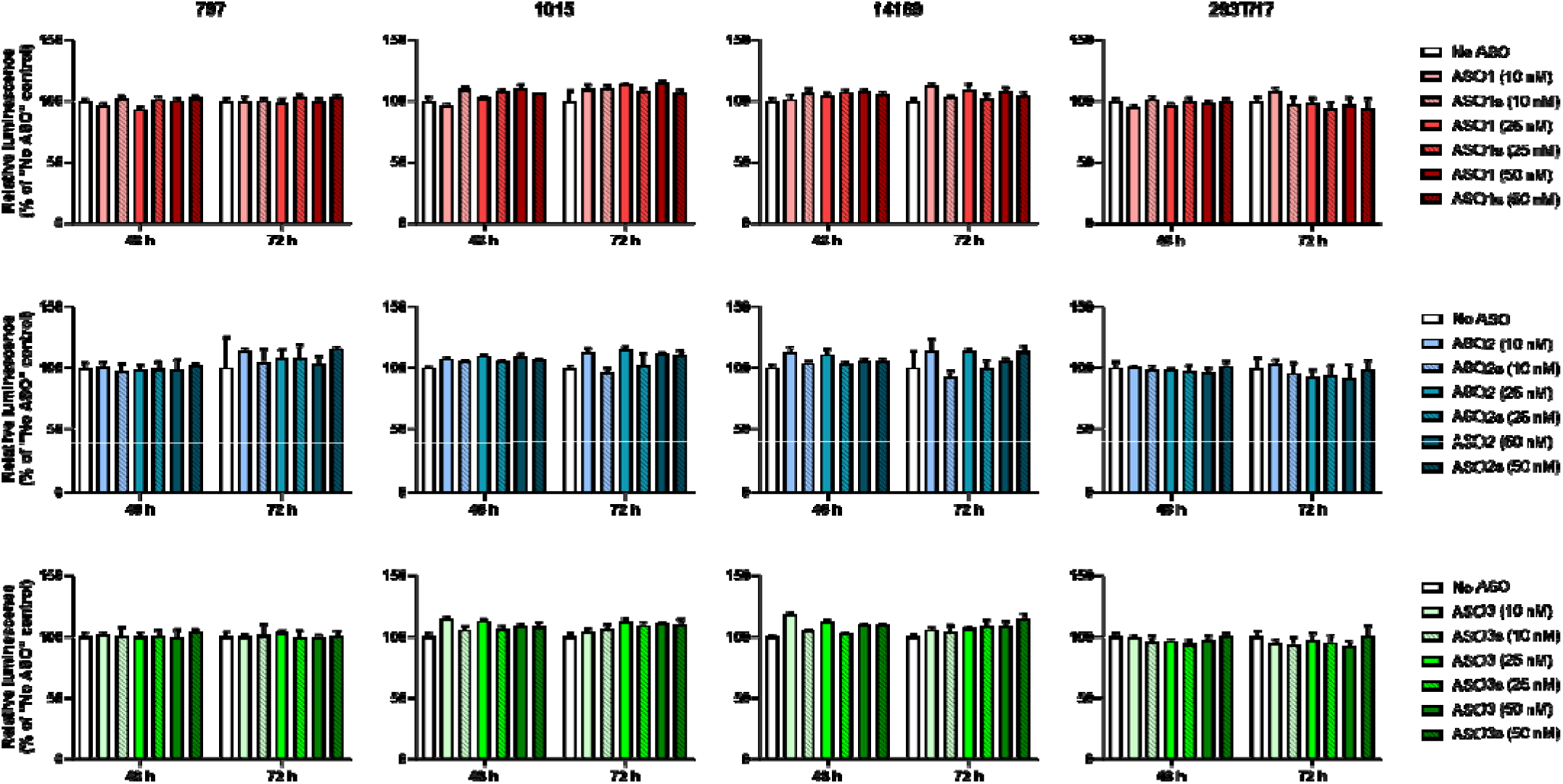
Three NUT carcinoma (797, 1015, 14169) and one non-NUT carcinoma (293T/17) cell lines were seeded in white 96-well plates at 15000 viable cells/well. The next day, antisense oligonucleotides targeting NUTM1 (ASO1, ASO2, ASO3) or scrambled counterparts containing the same respective nucleic acids in randomized order (ASO1s, ASO2s, ASO3s) were delivered in 10 µL Opti-MEM to reach the doses indicated. Either 48 or 72 h later, culture medium was replaced with DPBS and an equal volume (100 µL) of Promega CellTiter-Glo Reagent was added followed by shaking at 300 rpm for 5 min. Luminescence was recorded after an additional 5 min. Data are presented as means of three or six replicates for each measurement, depicted relative to respective plate’s negative control, with error bars indicating standard deviations.

**Figure 9.**
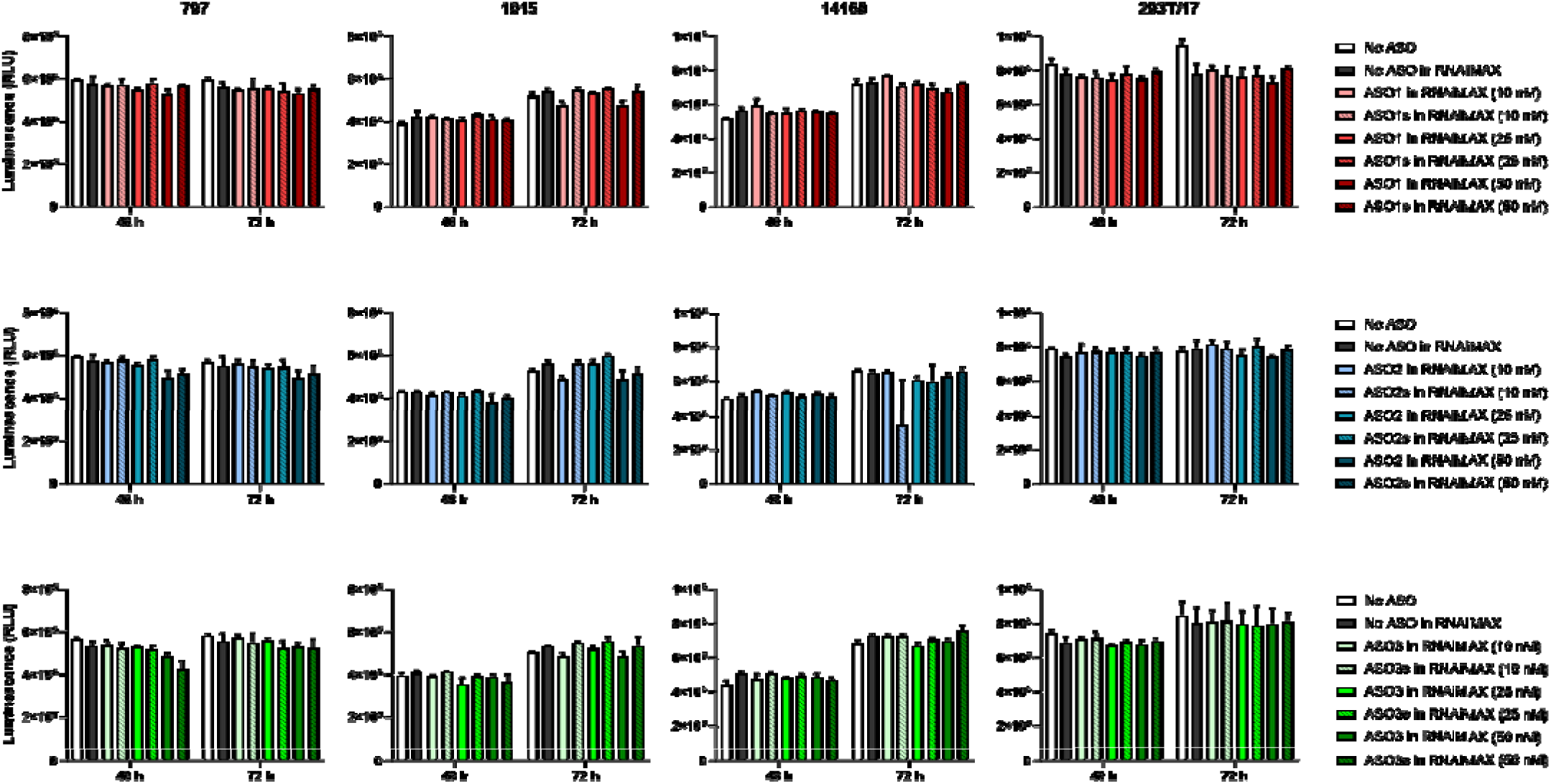
Three NUT carcinoma (797, 1015, 14169) and one non-NUT carcinoma (293T/17) cell lines were seeded in white 96-well plates at 15000 viable cells/well. The next day, antisense oligonucleotides targeting NUTM1 (ASO1, ASO2, ASO3) or scrambled counterparts containing the same respective nucleic acids in randomized order (ASO1s, ASO2s, ASO3s) were mixed with Lipofectamine RNAiMAX transfection reagent following manufacturer’s protocol and delivered at the doses indicated. Either 48 or 72 h later, culture medium was replaced with DPBS and an equal volume (100 µL) of Promega CellTiter-Glo Reagent was added followed by shaking at 300 rpm for 5 min. Luminescence was recorded after an additional 5 min. Data are presented as means of three replicates for each measurement, with error bars indicating standard deviations.

**Figure 10.**
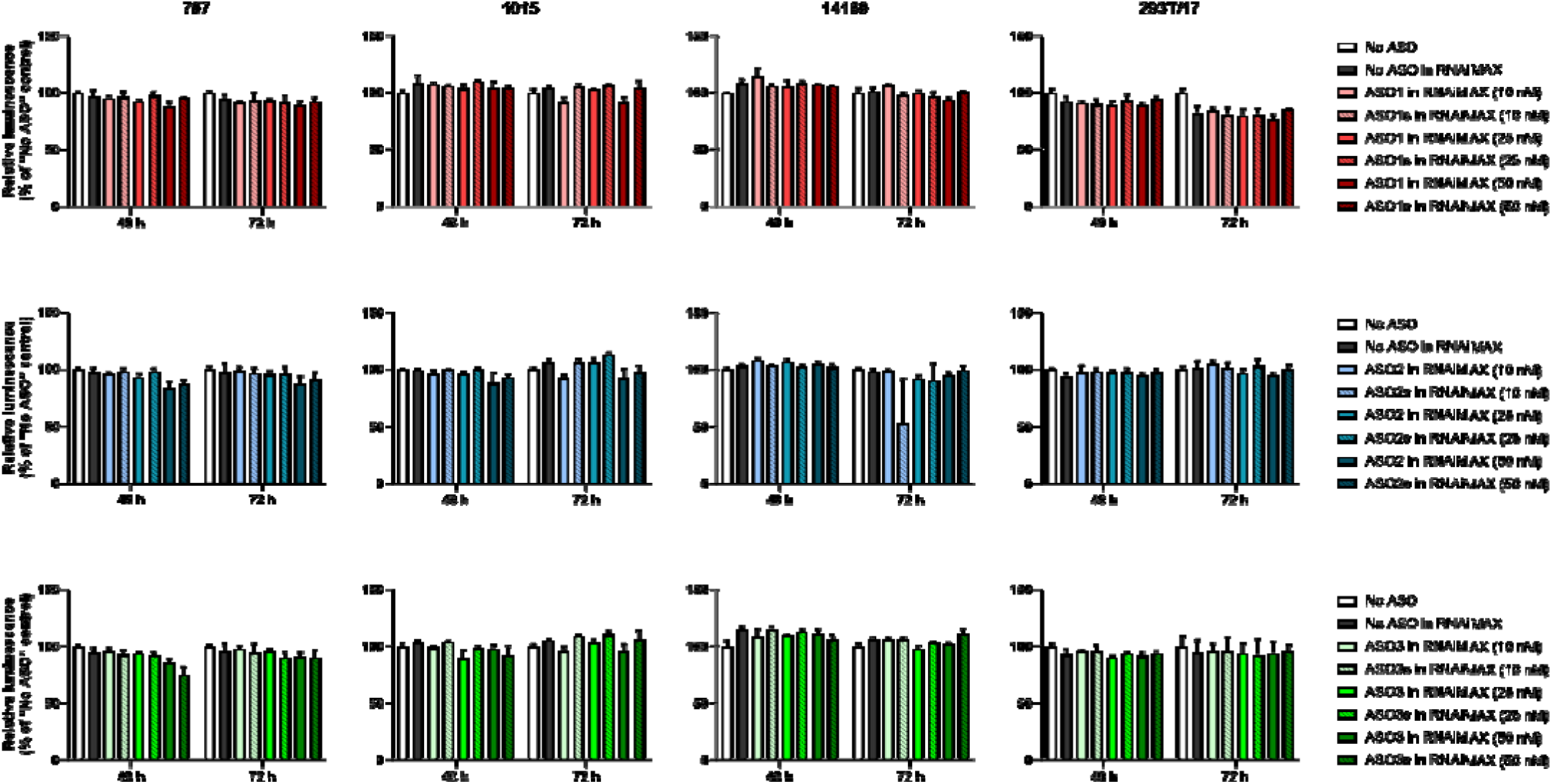
Three NUT carcinoma (797, 1015, 14169) and one non-NUT carcinoma (293T/17) cell lines were seeded in white 96-well plates at 15000 viable cells/well. The next day, antisense oligonucleotides targeting NUTM1 (ASO1, ASO2, ASO3) or scrambled counterparts containing the same respective nucleic acids in randomized order (ASO1s, ASO2s, ASO3s) were mixed with Lipofectamine RNAiMAX transfection reagent following manufacturer’s protocol and delivered at the doses indicated. Either 48 or 72 h later, culture medium was replaced with DPBS and an equal volume (100 µL) of Promega CellTiter-Glo Reagent was added followed by shaking at 300 rpm for 5 min. Luminescence was recorded after an additional 5 min. Data are presented as means of three replicates for each measurement, depicted relative to respective plate’s negative control, with error bars indicating standard deviations.

## Acknowledgments

We would like to thank Chris French’s Lab at Brigham and Women’s Hospital for providing us with the TC-797, 10-15 and 14169 NUT carcinoma cell lines. This work is funded by the Max Vincze Foundation and Ben Brown NUT Carcinoma Research Fund, both of whom we would like to thank for their support.

## Supplement: Experimental Protocol Document

### Prior to experiment week – Resuspend ASOs (completed on 1/26/24)

- Bring ASOs to room temperature
- Briefly spin each ASO tube (1 min at max speed/21130 rcf)
- Inside biosafety cabinet, resuspend each 200-nmol ASO tube to 100 µM stock concentration:

- Add 1 mL IDTE resuspension buffer, pH = 7.5 (IDT 11-01-02-02)
- Vortex and check that ASO visibly dissolved
- Quick spin contents to bottom of tube
- Add 1 mL IDTE resuspension buffer to bring total volume to 2 mL at 100 µM
- Invert several times to mix

Note: These IDT tubes can hold a max of approximately 2 mL with just enough space remaining to mix contents by inverting.

- Transfer 500 µL from each stock tube to new tubes to thaw for transfection
- Freeze all tubes at -20 °C

- Stock tubes stored in “JRA -20 freezer box” in Marsico freezer by tissue culture room
- Tubes for this experiment stored in tube rack in freezer in freezer hallway (will be moved to “JRA -20 freezer box” following transfection)

### Day 1 of experiment week – Plate cells (completed on 1/28/24)

- Remove media from cells in T75 flasks seeded at 5.5 x 10^6 cells/flask on 1/25/24 (with media changed on 1/26/24)

Note: Passage numbers of these cells had not been tracked prior to my handling of them.

- Add 10 mL cold Accutase and leave at room temperature until cells are dissociated
- Wash cells with 10 mL prewarmed media (spin at 300 x *g* for 5 min with brake = 1)
- Resuspend cells in 10 mL prewarmed media
- Use Countess to count:

- 797 – 5.62 x 10^6 cells/mL (99% viable) and 5.30 x 10^6 cells/mL (98% viable) = 5.38 x 10^6 viable cells/mL
- 1015 – 3.35 x 10^6 cells/mL (97% viable) and 3.75 x 10^6 cells/mL (98% viable) = 3.46 x 10^6 viable cells/mL
- 14169 – 3.48 x 10^6 cells/mL (98% viable) and 3.88 x 10^6 cells/mL (99% viable) = 3.68 x 10^6 viable cells/mL
- 293T/17 – 4.89 x 10^6 cells/mL (99% viable) and 5.00 x 10^6 cells/mL (99% viable) = 4.90 x 10^6 viable cells/mL
- Resuspend cells to 15000 <u>viable</u> cells/200 µL = 75000 <u>viable</u> cells/mL by bringing volume containing 3 x 10^6 cells to a total volume of 40 mL in each of two 50-mL conical tubes:

- 797 – 558 µL cell suspension + 39.442 mL media
- 1015 – 867 µL cell suspension + 39.133 mL media
- 14169 – 815 µL cell suspension + 39.185 mL media
- 293T/17 – 612 µL cell suspension + 39.388 mL media
- Plate 200 µL in respective wells of numbered white-walled, white bottom plates (Falcon 353296) with lids labeled

Note: Also numbered sides of plates in case lids get switched. Plated one clear 96-well plate to visually monitor growth and impact of some treatments.

- Placed in incubator at <u>3:03 pm</u>

### Day 2 of experiment week – Transfect cells with ASOs ∼24 h post-plating (completed 1/29/24)

- Bring Lipofectamine RNAiMAX to room temperature
- Prepare each ASO for gymnotic delivery:

- Dilute each ASO from 100 µM (stock concentration) to 1 µM by adding 10 µL to 990 µL Opti-MEM; will add 10 µL of this to respective 50-nM wells
- Dilute 1-µM ASO solutions to 500 nM by adding 200 µL to 200 µL Opti-MEM; will add 10 µL of this to respective 25-nM wells
- Dilute 1-µM ASO solutions to 200 nM by adding 200 µL to 800 µL Opti-MEM; will add 10 µL of this to respective 10-nM wells
- One ASO dose at a time, deliver each ASO to plates 1-6 as described above such that only two plates are ever out of the incubator at the same time

- Time that this step began: <u>2:55 pm</u>

Note: These volumes were adequate to use a P20 multichannel with a 3-well reservoir.

- Prepare each ASO for delivery with Lipofectamine RNAiMAX:

- 50-nM wells – no dilution necessary (will use stock concentration of 100 µM)
- 25-nM wells – dilute each ASO from 100 µM (stock concentration) to 50 µM by adding 5 µL to 5 µL Opti-MEM
- 10-nM wells – dilute each ASO from 100 µM (stock concentration) to 20 µM by adding 5 µL to 20 µL Opti-MEM
- Prep and label tubes for delivery with Lipofectamine RNAiMAX:

- Add 141.5 µL Opti-MEM to each of 18 tubes (will receive RNAiMAX)
- Add 196 µL Opti-MEM to each of 18 tubes (will receive ASO)
- One ASO at a time, prepare transfection mix and deliver to plates 7-12 as described below such that only two plates are ever out of the incubator at the same time:

- Add 8.5 µL RNAiMAX to tube already containing 141.5 µL Opti-MEM
- Add 4 µL ASO to tube already containing 196 µL Opt-MEM then transfer 150 µL of this to previous tube (containing RNAiMAX plus Opti-MEM) and mix
- Wait 5 min at room temperature
- Add 10 µL to respective wells

Note: These volumes were just adequate to use a P20 multichannel with a 3-well reservoir.

- Prepare and deliver RNAiMAX to “No ASO in RNAiMAX” wells:

- Add 354 µL Opti-MEM to a tube then add 21.2 µL RNAiMAX then add 375 µL Opti-MEM and mix
- Wait 5 min at room temperature
- Add 10 µL to respective wells (row D of plates 7-12)
- Return all plates to the incubator

- Time that this step ended: <u>6:30 pm</u>

### Day 4 of experiment week – Read plates ∼48 h after transfection (completed 1/31/24)

- Bring reconstituted CellTiter-Glo Reagent to room temperature in the dark

- Mix reagent from two kits (lot # 0000181000 and lot # 0000169392) 1:1 after thawing
- Remove odd-numbered plates from incubator (six total) and transport to Marsico

Note: Also checked clear 96-well plate at this time. 797 and 293T/17 cells appeared nearly 100% confluent with 1015 and 14169 cells slightly less confluent. Media color was not noticeably different in any wells nor were any cells visibly impacted by treatments on this plate (though only a few treatments were plated here).

- Staggering plates, dump contents and replace with 100 µL room temperature DPBS
- Add 100 µL room temperature CellTiter-Glo Reagent to each well
- Shake each plate for 5 min at 300 rpm then wait an additional 5 min at room temperature before using plate reader to record luminescence

### Day 5 of experiment week – Read plates ∼72 h after transfection (completed 2/1/24)

- Bring reconstituted CellTiter-Glo Reagent to room temperature

- Mix reagent from two kits (lot # 0000181000 and lot # 0000169392) 1:1 after thawing
- Remove even-numbered plates from incubator (six total) and transport to Marsico Note: Also checked clear 96-well plate at this time. 797 and 293T/17 cells appeared completely confluent with media color more orange in these wells (coloring also followed this pattern on even-numbered white plates). No cells were visibly impacted by treatments on this plate (though only a few treatments were plated here).

- Staggering plates, dump contents and replace with 100 µL room temperature DPBS
- Add 100 µL room temperature CellTiter-Glo Reagent to each well
- Shake each plate for 5 min at 300 rpm then wait an additional 5 min at room temperature before using plate reader to record luminescence

